# Computing the inducibility of B cell lineages under a context-dependent model of affinity maturation: Applications to sequential vaccine design

**DOI:** 10.1101/2023.10.13.562156

**Authors:** Joseph Mathews, Elizabeth Van Itallie, Kevin Wiehe, Scott C. Schmidler

## Abstract

A key challenge in B cell lineage-based vaccine design is understanding the *inducibility* of target neutralizing antibodies. We approach this problem through the use of detailed stochastic modeling of the somatic hypermutation process that occurs during affinity maturation. Under such a model, sequence mutation rates are *context-dependent*, rendering standard probability calculations for sequence evolution intractable. We develop an algorithmic approach to rapid, accurate approximation of key marginal sequence likelihoods required to inform modern sequential vaccine design strategies. These calculated probabilities are used to define an inducibility index for selecting among potential targets for immunogen design. We apply this approach to the problem of choosing targets for the design of boosting immunogens aimed at elicitation of the HIV broadly-neutralizing antibody DH270min11.

**Author summary:** Vaccine effectiveness relies on inducing the immune system to generate protective antibodies. Because antibodies are generated by random processes coupled to positive selection, the ability to induce certain rare but desirable antibodies can be limited by the inherent probability of occurrence. We use computational modeling to estimate the probability of antibody occurrence, and demonstrate the use of these estimates in designing vaccine regimens which maximize the probability of induction of broadly neutralizing antibodies.

## 1 Introduction

Vaccination aims to induce antibodies – the secreted versions of B cell receptors – that can neutralize pathogens and thus provide protection against infection. In natural infection, B cells evolve their antigen receptors to specifically recognize pathogens through progressive rounds of mutation and selection based on affinity to antigen.

However, rapidly mutating viruses such as HIV, influenza and SARS-CoV-2 can escape the B cell response through viral diversification. In such cases, it is of great interest to develop vaccination strategies to elicit *broadly neutralizing* antibodies (bnAbs). However, bnAbs are rarely elicited in infection due to a variety of factors [1]. For example, HIV bnAbs typically originate from B cells with low precursor frequencies in the human B cell receptor repertoire [2, 3]. They also typically have high numbers of mutations, some of which may be essential for broad neutralization but be made infrequently by activation-induced cytidine deaminase (AID), the enzyme responsible for mutating the B cell receptor [4]. Acquisition of these improbable mutations is a key bottleneck for bnAb induction [4, 5]. In these situations, traditional vaccine design strategies have proven ineffective at eliciting bnAbs. This has motivated the development of advanced vaccine design strategies which aim to use known bnAbs as templates to design immunogens that can direct B cell evolution towards their induction [2, 6–9].

One promising HIV vaccine design approach, called *sequential prime-boosting*, aims to first induce low-frequency bnAb B cell precursors with a *priming* immunogen, and then mature those clonal lineages with one or more additional, distinct *boosting* immunogens. A key step of this approach is the inference of a reconstructed B cell receptor (BCR) sequence of the bnAb precursor, referred to as the unmutated common ancestor (UCA), from observed, clonally-related sequences [6, 10]. Recent progress has been made on the first component of this sequential prime-boost approach through the design of priming immunogens that can initiate precursors of bnAb B cell clonal lineages [11]. However, the design of boosting immunogens that can select for sets of required, improbable mutations in the initiated bnAb B cell clonal lineages remains a central challenge [2].

Current experimental approaches to develop sequential prime boost vaccine regimens are highly labor- and time-intensive, relying on cycles of immunogen design, immunization of animal models, and B cell receptor repertoire sequencing to assess whether desired mutations were elicited [5, 8, 12]. An alternative approach, which we adopt here, is the use of computational models to accelerate the design process.

Specifically, we consider the use of stochastic models of the somatic hypermutation process to better inform the design of immunogens and development of vaccine regimens to efficiently guide a maturing bnAb response by vaccination.

Critical to this approach is the recognition that fixation of mutations is the product of two separate forces: mutation and selection. While vaccination with carefully chosen immunogens can introduce targeted selection, it is hypothesized that one of the primary difficulties in eliciting bnAbs (in HIV, say) is the low frequency with which essential mutations occur [4]. If this frequency is sufficiently low, there are unlikely to be any instances of the mutation in the B cell population on which this selection can act. The use of stochastic models of the somatic hypermutation process to understand this frequency is therefore critical. A key step in formalizing this problem is answering the question: given an ancestral B cell receptor sequence **x** within a B cell clonal lineage, what is the probability

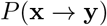

of obtaining a specified target sequence **y** along the lineage under a realistic model of affinity maturation. However, because mutation of B cell receptors is sequence context-dependent due to biases in mutational targeting and substitution by AID, mutations do not arise independently during B cell evolution. This context dependence raises the key technical challenge to be addressed in this paper, namely that the context-dependent nature of somatic hypermutation makes standard approaches for computing *P* (**x** → **y**) under molecular evolution models intractable.

### Challenges in Sequential Prime Boost Vaccine Design

Practical limitations on the number of boosting immunogens that can be administered mean that optimizing the sets of mutations to be selected by each immunogen is an important consideration in the design of a sequential prime-boosting vaccine regimen. Due to the sequence context-dependence of somatic hypermutation, the order in which the immunogens are administered and select for desired mutations within a vaccine regimen will also affect the probability of eliciting a bnAb response. Additionally, since it is not known *a priori* what the success of the chosen immunogens will be, the vaccine designer must make choices about which mutations to target first without information about the complete vaccine regimen. The ability to calculate *P* (**x** → **y**) enables us to compute more generally *P* (**x**^(1)^ → **x**^(2)^ → … **x**^(*p−*1)^ → **y**) for any ordered set of intermediate sequences **x**^(2)^, …, **x**^(*p−*1)^, a key step for such design choices. The ability to accurately calculate these full-length BCR sequence evolution probabilities, and to rank BCR sequences by their *inducibility*, has a number of practical applications in the sequential vaccine design process. We outline three common scenarios encountered in designing sequential prime-boost vaccine strategies where such calculations can be used to inform the design of specific immunogens:

***(Design scenario I)*** In *ab initio* lineage-based vaccine design, the first step is to design a priming immunogen that optimally engages bnAb precursor B cells and maximizes the probability that B cells will evolve along bnAb maturation pathways. We assume that the priming immunogen already binds with high affinity to the unmutated precursor bnAb B cells. Given that a limited number of mutations can be selected by any one immunogen, the challenge is to identify the set of mutations to target for selection first, in order to give the B cells the highest chance of success to eventually mature into bnAbs.

***(Design scenario II)*** At intermediate stages of the design process, the vaccine designer has developed an initial set of immunogens in a vaccine regimen and has evaluated the B cell response to this partial regimen by sequencing the antibody repertoire of immunized bnAb UCA knock-in mice [5, 8, 12–14]. By measuring the frequency of occurrence of targeted mutations in immunized knock-in mice, the vaccine designer can determine which of the mutations necessary for broad neutralization are induced by immunization with the regimen. Often the regimen is partially successful, in that a subset of the necessary mutations are selected, leaving others still in need of selection by addition of subsequent immunogens to the regimen. During this iterative process, the probability of evolving the target bnAb *conditional on* the observed intermediate(s) can be used to identify the set of mutations to target for selection with the next boosting immunogen, in order to maximize the probability of bnAb induction.

***(Design scenario III)*** To induce bnAb maturation, immunogens are designed to bind to antibodies with desired mutations within the targeted bnAb lineage. While *in vitro* binding assay data can be predictive of mutation selection, ultimately the ability of the immunogen to select for specific mutations must be measured *in vivo* in animal model immunization studies. During this intermediate stage of vaccine design, experimentally observed mutation occurrence frequencies are available from immunized UCA knock-in mice. The vaccine designer may then compare mutation probabilities in the absence of selection (calculated under the model) with the experimentally observed mutation frequencies, in order to identify mutations that are selected for or against by the current regimen.

### Organization of this paper

The remainder of this paper is organized as follows. In Section 2 we develop a fast and accurate approximation algorithm for the problem of calculating full-length sequence transition probabilities *P* (**x** → **y**) under a stochastic, context-dependent model of somatic hypermutation. In Section 3.1, we evaluate the accuracy and speed of this approach on problems of varying difficulty. In Section 3.2, we demonstrate the use of this approach to calculate key quantities required in the above design scenarios to enable rational selection of immunogens for a sequential prime-boosting strategy.

### 2 Methods

#### 2.1 Background and Motivation

The ARMADiLLO model [4] is a recently developed model for forward simulation of the somatic hypermutation process in affinity maturation. The simulation procedure uses a set of mutation rates and base frequencies derived from NGS data [15]. Unlike the continuous-time Markov chain (CTMC) models common in molecular evolution, sites in the ARMADiLLO model evolve in discrete jumps, a process that can be viewed as the skeleton of a time-inhomogeneous CTMC. However, a critical aspect of the somatic hypermutation model encoded in ARMADiLLO is the sequence *context-dependence* of mutation and substitution rates, arising from the sequence targeting preferences of the AID enzyme. Just as in dependent-site models of sequence evolution [16–18], calculating *p*(**y** | **x**) under the ARMADiLLO model is computationally difficult due to the dependence among sites which precludes the use of Felsenstein’s pruning algorithm [19]. Although forward simulation of the ARMADiLLO model has been used to estimate the probability of individual mutations arising by chance [4], this approach becomes prohibitively expensive when trying to calculate *p*(**y** | **x**) for a specific, full sequence **y**. In Section 2.3, we develop an importance sampling algorithm for tractably approximating *p*(**y** | **x**). Our approach samples mutation orderings (trajectories) of observed mutations rather than entire path histories as in previous dependent-site models, and handles the multimodality that can plague other sampling approaches.

Section 3 demonstrates that our method can be used to easily approximate a large number of such transition probabilities quickly.

#### 2.2 The ARMADiLLO Model of Somatic Hypermutation

Let **x** = (*x*_1_, *x*_2_, …, *x*_*n*_) and **y** = (*y*_1_, *y*_2_, …, *y*_*n*_) denote two nucleotide sequences. Let 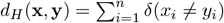 denote the Hamming distance between **x** and **y** and let 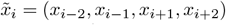 for *i* ∈ {1, …, *n*} denote the two-nearest-neighbor context of site *x*_*i*_. (For *i* = 1 and *i* = *n*, we assume that the two left and right flanking nucleotides, respectively, are fixed). Each site *x*_*i*_ is assigned a mutability score 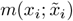 and a set of substitution probabilities 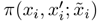, where 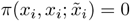 and 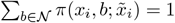 for 𝒩= {*A, G, C, T*}. The ARMADiLLO algorithm is given in Algorithm 1. Note that the transition probability under ARMADiLLO is defined by a probability matrix *Q*^(*i*)^ with entries

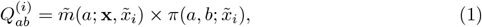

where

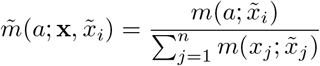

denotes the normalized mutability score. The factors 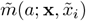 and 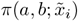 appearing in (1) correspond to the probability of selecting site *x*_*i*_ for mutation and transitioning to nucleotide base *b*, respectively. For information on how the mutability scores and transition probabilities are estimated, see [15].

##### Algorithm 1 ARMADiLLO

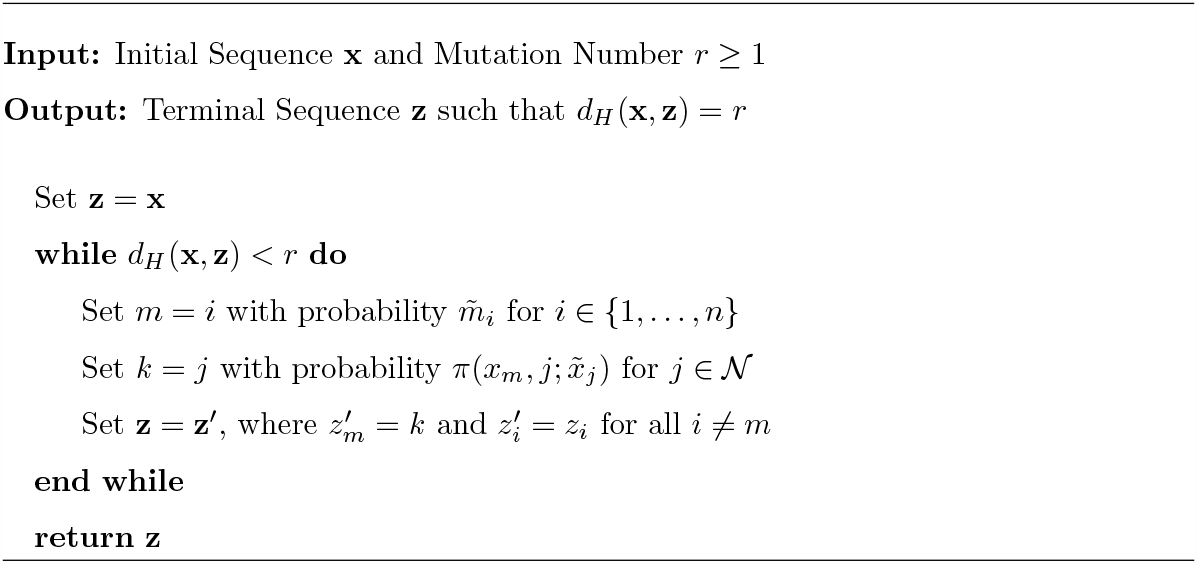

Algorithm 1 was used [4] to estimate the marginal probability *p*(*y*_*i*_ | **x**) of a codon at a given location in **x** transitioning to a target amino acid by simulating many terminal nucleotide sequences **y**^(1)^, …, **y**^(*M*)^ and calculating the proportion 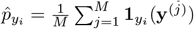 of times that the target amino acid occurs at the desired location. Because the emphasis is on the marginal probability *p*(*y*_*i*_ | **x**) at a single site, a reliable estimate of this probability appears to be obtainable with a feasible number of simulations. However, as noted this approach does not scale to computation of *p*(**y** | **x**) for full-length sequences **y**. Instead, we develop an efficient importance sampling algorithm for evaluating full-length sequence probabilities of this form.

#### 2.3 Estimation using Importance Sampling

Given two sequences **x** and **y** such that *r* = *d*_*H*_ (**x, y**), we wish to approximate *p*(**y** | **x**) under Algorithm 1. To do this, we sample orderings in which the *r* mutations occur in the transition from **x** to **y**. More formally, let 𝒮 = (*s*_1_, …, *s*_*r*_) for *s*_*j*_ ∈ {1, …, *n*} be the set of sites at which the two sequences differ. Let *S*_*r*_ be the set of all permutations of the elements of 𝒮. The goal is to approximate

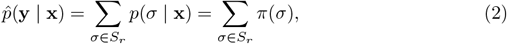

where

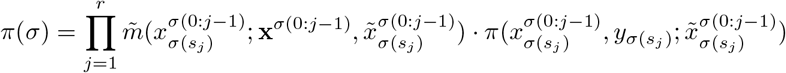

and 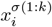 denotes, for a given *σ* ∈ *S*_*r*_, the value of the *i*th nucleotide after *k* updates to **x** according to the permutation *σ*. That is

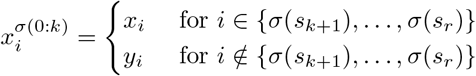

(so e.g. 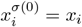, and 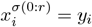). We refer to *σ* ∈ *S*_*r*_ as a ‘path’ from **x** to **y**.

Let 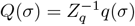 where 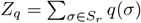 denote a distribution on *S*_*r*_ (the instrumental distribution) which can be easily sampled. Then the corresponding importance sampling estimator for (2) is

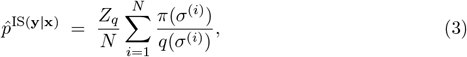

for 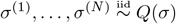. The performance of this importance sampling algorithm depends strongly on the choice of *q*, with max_*σ*_ *π/q* controlling Var 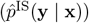. Because *π* is unnormalized here, we require *Z*_*q*_ in order to recover 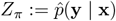. Consequently, we need to choose *Q* complex enough to approximate *π* reasonably well but simple enough so that *Z*_*q*_ can be feasibly computed.

Recall that under the ARMADiLLO model the mutability score 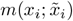 for a site *x*_*i*_ depends only on its two-nearest-neighbor context 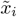. However, *x*_*i*_ mutates to *a* ∈ 𝒩 with probability equal to its *normalized* mutability score 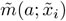. This makes the normalization *Z*_*π*_ for the ARMADiLLO path distribution intractable because the product of normalization terms

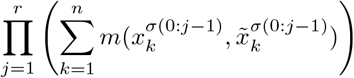

appearing in *π*(*σ*) induces long-range dependence among sites that do not have overlapping pentamers, and consequently the probability that *x*_*i*_ mutates depends on *all* sites *x*_1:*n*_ := (*x*_1_, …, *x*_*n*_). This suggests replacing 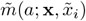 with 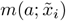 in *π*(*σ*) to make *Z*_*π*_ tractable; we use this form as our instrumental distribution *Q*. In particular, we set

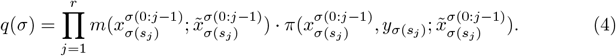

Since the context 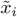 plays the most important role in determining the mutation probability for *x*_*i*_, this provides an instrumental distribution *Q* that closely approximates *π* while also yielding a tractable normalizing constant *Z*_*q*_.

To see that *Z*_*q*_ is tractable, call two subsets **s, s**^*′*^⊂𝒮 *separated* if for any *s* ∈ **s** and *s*^*′*^∈ **s**^*′*^ we have |*s* − *s*^*′*^| *>* 2. Under *q*, the probability of a permutation is invariant with respect to the re-ordering of mutations belonging to different separated sets. More formally, let (**s**_1_, …, **s**_*k*_) be a partition of S into separated sets. Let *r*_*j*_ = |**s**_*j*_| and let 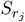 be the set of all permutations of the elements of 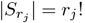 with 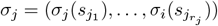 denoting a corresponding permutation 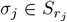. Let

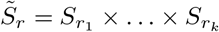

be the product group consisting of all *k*-tuples (*σ*_1_, …, *σ*_*k*_) with 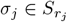 and note that 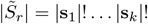. Consider the equivalence class of 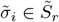

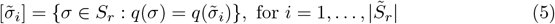

Provided 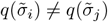 for *i* ?= *j* (this will generally be the case), then the equivalence classes (5) partition *S* into 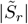 sets and 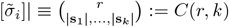 for *all* 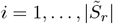 (see Example 1 below). Hence,

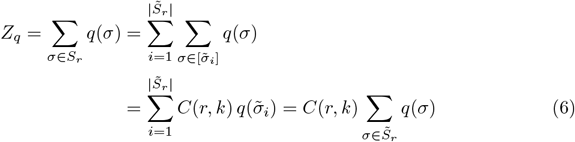

Putting it all together, we have by (2), (3), and (6) the estimator

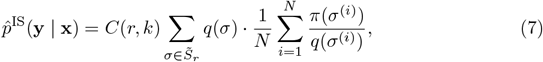

for 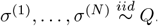. Then 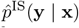 is an unbiased estimator of 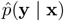.

Sampling from *q* proceeds by enumerating all 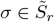 and evaluating (4). *Z*_*q*_ is then obtained by summing and multiplying by *C*(*r, k*) as in (6). We can then sample index *I* with probability *q*(*σ*_*i*_)*/Z*_*q*_ for 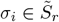 corresponding to one of the equivalence classes 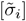. Finally, we sample uniformly from the equivalence class 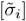 to obtain the desired sample from *q*.

Modes of *π* are obtained at any permutations which maintain ‘optimal’ orderings on the subsets **s**_1_, …, **s**_*k*_. We can identify these optimal orderings by computing 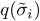 for all 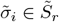. Since *q* is constant over 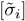, there are at least *C*(*r, k*) such modes. Using *Q* as our instrumental distribution allows us to sample from this multiplicity of high probability regions efficiently. In contrast, Markov chain Monte Carlo sampling - which has been applied to other context-dependent modeling problems [16, 17] in molecular evolution - can often suffer from slow mixing in the face of such multimodality.

**Example 1**. Suppose **x** and **y** are two sequences of length *n* = 10 differing at sites *S* = (3, 4, 7, 8). Let **s**_1_ = (3, 4), **s**_2_ = (7, 8) be a partition of *S* into separated sets. Then

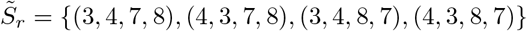

and *S*_*r*_ is the set of all permutations of (3, 4, 7, 8). Consider element 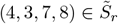.

There are 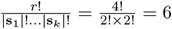 permutations which yield the same value as (4, 3, 7, 8) under (4). These are (4, 3, 7, 8), (4, 7, 3, 8), (4, 7, 8, 3), (7, 8, 4, 3), (7, 4, 8, 3), and (7, 4, 3, 8). In all cases, 4 and 7 appear before 3 and 8, respectively, and therefore ‘respect the orderings’ for *σ*_1_(**s**_1_) = (4, 3) and *σ*_2_(**s**_2_) = (7, 8).

### 3 Results

We begin with a simulation study to evaluate the performance of our approach, followed by the application to design calculations for HIV-I immunogen design.

#### 3.1 Simulation Study

We first evaluate the approximation algorithm of Section 2 on a set of example sequences where the true transition probabilities can be calculated exactly. Test sequences are chosen to span a range of values of both the largest separated set *r*^*⋆*^ := max_*j*_ *r*_*j*_ and the quantity *α* := *r*^2^*/n* which quantifies the number of mutations *r* relative to the size of the sequence *n*. (We expect that the variance of our estimator increases with *α*.) This set of test sequences demonstrates the performance of the algorithm as the complexity of the problem varies. We investigate the performance of the estimator (7) as a function of *r*^*⋆*^ and *α*, measured in terms of both coefficient of variation and effective sample size (ESS):

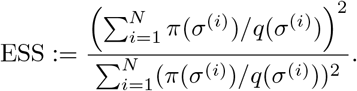

In all test sequences, the number of mutations is kept low (*r* = 10) so that the exact value of the transition probability can be computed for comparison.

For each test sequence, the algorithm was run 1000 times using a sample size *N* = 1000 in each run. Table 1 shows the results, including the exact value, and the mean and relative standard deviation of the 1000 estimates. Across all values of *r*^*⋆*^ and *α* the transition probability is estimated accurately up to at least the second significant digit. The coefficient of variation decays with *α* as expected, and appears to be unaffected by *r*^*⋆*^. The most difficult case is when *α* and *r*^*⋆*^ are large (Table 1, first row), whereas the ideal case is when both of these quantities are small (last row), and we see this difficulty reflected in the average effective sample size.

**Table 1.** Results for estimating transition probabilities on test sequences. 1000 runs of the algorithm were performed for each sequence, each using *N* = 1000. Shown are exact calculations (True prob.), mean estimate over the 1000 replicates, mean effective sample size, and standard deviation of estimates relative to the mean (CoV). We see that probabilities are estimated accurately across all values of *r*^*⋆*^ and *α*, despite the problems getting more challenging (lower ESS, higher CoV) when *α* and *r*^*⋆*^ are large.

| Test seq. | $r^*$ | $\alpha$ | True prob. | Avg. Est. | Avg. ESS | C.o.V. |
| --- | --- | --- | --- | --- | --- | --- |
| 1 | 6 | 2 | $3.362 \times 10^{-15}$ | $3.360 \times 10^{-15}$ | 397 | 0.027 |
| 2 | 3 | 2 | $4.695 \times 10^{-14}$ | $4.691 \times 10^{-14}$ | 475 | 0.021 |
| 3 | 0 | 2 | $6.703 \times 10^{-18}$ | $6.696 \times 10^{-18}$ | 679 | 0.022 |
| 4 | 6 | 1.5 | $1.178 \times 10^{-15}$ | $1.178 \times 10^{-15}$ | 570 | 0.014 |
| 5 | 3 | 1.5 | $1.798 \times 10^{-18}$ | $1.796 \times 10^{-18}$ | 714 | 0.012 |
| 6 | 0 | 1.5 | $2.833 \times 10^{-20}$ | $2.833 \times 10^{-20}$ | 864 | 0.012 |
| 7 | 6 | 0.5 | $7.198 \times 10^{-23}$ | $7.199 \times 10^{-23}$ | 629 | 0.004 |
| 8 | 3 | 0.5 | $7.050 \times 10^{-22}$ | $7.050 \times 10^{-22}$ | 682 | 0.005 |
| 9 | 0 | 0.5 | $3.189 \times 10^{-24}$ | $3.188 \times 10^{-24}$ | 943 | 0.008 |

#### 3.2 Application to B-Cell Evolution

We now return to the problem of sequential immunogen design and the scenarios described in Section 1 encountered by vaccine designers in which the transition probabilities between the UCA sequence and target mature antibody sequences can be used to inform the design of specific immunogens.

For example, Fig 1 shows (see also Table 3 below) that the order of mutation selection in a clonal lineage can significantly affect the probability of obtaining specific bnAb maturation pathways. Since multiple immunogens may be required to direct the evolution of B cells when a large number of mutations must be acquired for bnAb activity [13, 20], the ordering of prime and boosting immunogens - each selecting distinct sets of mutations - within a lineage-based vaccine regimen may critically affect the probability of successful bnAb elicitation. It is therefore of great interest to determine a mutation ordering that maximizes this elicitation probability for a specified target bnAb. This in turn can be used to develop a sequence of immunogens to target bnAb mutations in the most probable order of acquisition.

**Figure 1.**
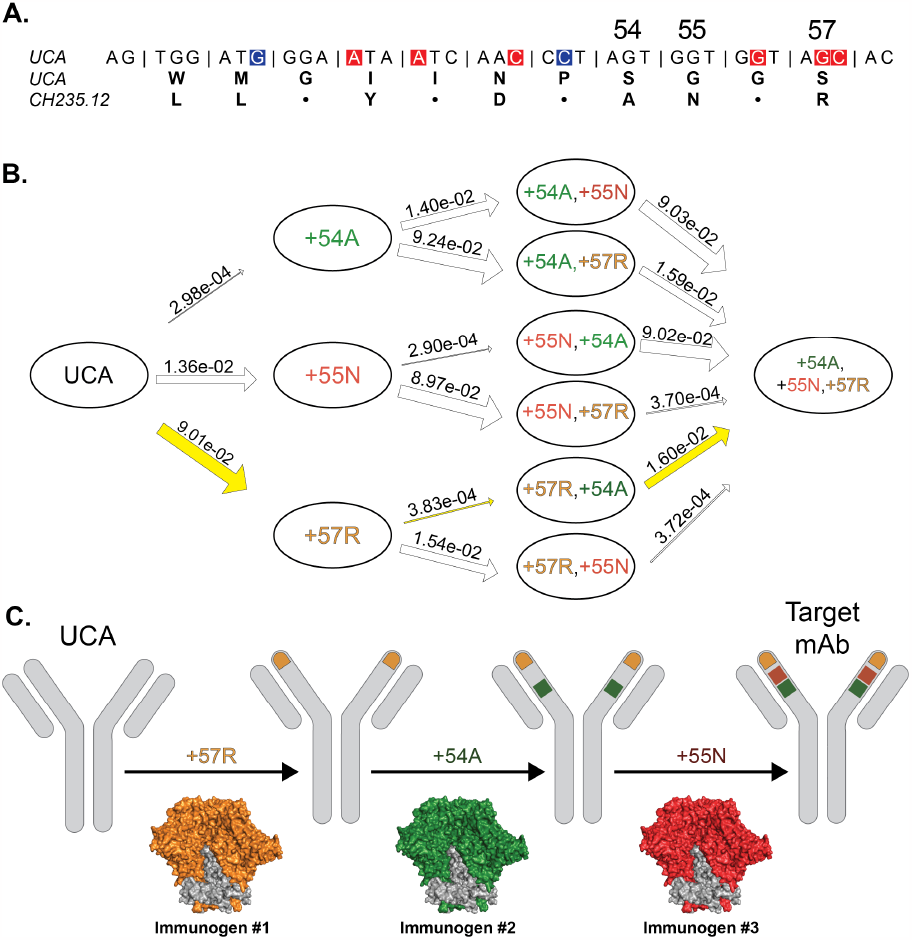
The use of estimated transition probabilities to inform sequential prime-boost vaccine design. **A)** CDRH2 sequence of the HIV bnAb CH235.12 inferred UCA. AID hot spots (high mutability) are highlighted in red and cold spots (low mutability) highlighted in blue, with amino acid alignment of the UCA and CH235.12 mature sequence shown below. Dots represent sequence matches and denote unmutated amino acid positions. **B)** Graph of amino acid transitions for a subset of three mutations at sites 54, 55, and 57. Arrow widths are proportional to estimated transition probabilities (shown). Based on the highest probability full path (yellow), **C)** the vaccine designer can choose the order of immunogens in a sequential prime-boost vaccine regimen to maximize the probability of induction of the three targeted mutations.

##### 3.2.1 Transition probabilities for protein sequences

In calculating maturation pathway probabilities, consideration of both nucleotide and the amino acid sequences is critical. The UCA sequence is the result of recombination of germline-encoded gene segments and addition of non-templated nucleotides at the gene segment junctions, providing a defined starting nucleotide sequence. However, selection acts upon the BCR at the protein level, where binding to the immunogen is determined by the amino acid sequence, and thus all nucleotide sequences giving rise to the target bnAb amino acid sequence must be accounted for. Because AID acts at the nucleotide level, the ARMADiLLO model describes transitions between nucleotide sequences.

Marginalization is then required to obtain probabilities of amino acid sequences. In what follows then, we compute the probability that a known unmutated common ancestor (UCA) nucleotide sequence transitions to a target *amino acid* sequence. For purposes of vaccine design, it will also be of interest to consider “intermediate” amino acid sequences along potential mutation pathways, as potential targets for sequential immunogen design. We formalize these calculations below.

Let *a*(**y**) denote the amino acid sequence arising from a nucleotide sequence **y** and *A*(**y**) = {**z** : *a*(**z**) = *a*(**y**)} the equivalence class of nucleotide sequences giving rise to the same amino acid sequence. Denote by *a*_*i*_(**y**) the *i*th amino acid in *a*(**y**), and let *c*_*i*_(**y**) denote the number of codons that encode *a*_*i*_(**y**). We estimate transition probabilities from an inferred UCA (initial) nucleotide sequence **x** to a *any* terminal nucleotide sequence **z** ∈ *A*(**y**), where **y** is an observed bnAb (target nucleotide sequence).

Let **m** = {*m*_1_, …, *m*_*k*_} for 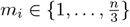 be the set of amino acid sites where *a*(**x**) and *a*(**y**) disagree and **m**_1_, …, **m**_*p*_ be all 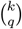 subsets of **m** of size *q*, with elements 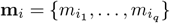. Let 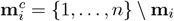 and define

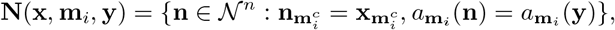

where 𝒩 denotes the set of nucleotides. (The condition 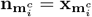 is a slight abuse of notation, indicating that all the codons outside of the set **m**_*i*_ are equal.) So **N**(**x, m**_*i*_, **y**) is the set of nucleotide sequences of length *n* which match **x** in all positions 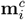, and which map to the same amino acids as **y** in positions **m**_*i*_. (**N** is a set due to the redundancy of the genetic code.)

Denote by **x**^*ij*^ the *j*th element of **N**(**x, m**_*i*_, **y**). So **x**^*ij*^ is an intermediate nucleotide sequence on a trajectory from **x** to some sequence in *A*(**y**) which is equal to **x** outside of (the codons indexed by) **m**_*i*_ and matches *a*(**y**) at all sites **m**_*i*_ (i.e.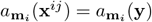).

The superscript *j* indexes the possible combinations of codons giving rise to 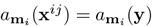, of which there are 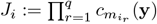 for *i* = 1, …, *p*.

Let **z** be an end-of-trajectory sequence for **x**^*ij*^, meaning that **z** ∈ *A*(**y**) and **z** differs from **x**^*ij*^ only at nucleotide positions corresponding to amino acid sites **m** \ **m**_*i*_ of the mutations not yet acquired by **x**^*ij*^. (That is, 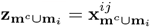.) We denote the set of all such **z** as *Z*(**x, y**, *i, j*) = *A*(**y**) ∩ **N**(**x**^*ij*^, **m** \ **m**_*i*_, **y**).

We estimate the transition probability of **x**_*i,j*_ to each **z** ∈ *Z*(**x, y**, *i, j*) by assuming exactly *r* = *d*_*H*_ (**x**^*ij*^, **z**) mutational events occur (i.e. unmutated nucleotide positions remain fixed, and no reversions). Starting from intermediate **x**^*ij*^ for *i* = 1, …, *p* and *j* = 1, …, *J*_*i*_, this yields transition probability

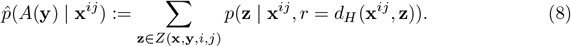

Similarly, the probability of obtaining the initial amino acid mutations in **m**_*i*_ is given by

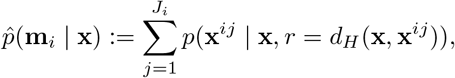

where again the conditioning indicates that exactly *r* mutations occur in the transition from **x** to **x**^*ij*^, with all other nucleotide positions remaining fixed. Finally, we calculate the joint probability of first obtaining initial amino acid mutations **m**_*i*_ on the way to obtaining the full set of mutations **m** as the joint probability:

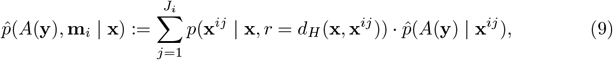

Conditioning on the number of mutational events that occur and assuming unmutated nucleotide positions remain fixed in our estimates amounts to ignoring synonymous mutations and multiple-mutation reversions. We make these assumptions for computational tractability. Indeed, if 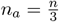 is the length of *a*(**y**), then the number of terminal sequences that give rise to *a*(**y**) can grow exponentially in *n*_*a*_ via synonymous mutations due to the redundancy in the genetic code.

##### 3.2.2 Inducibility of a minimal set of critical mutations in an HIV bnAb

We applied this approach to study the inducibility of critical mutations in DH270.6, an HIV bnAb. Here the target sequence **y** is the heavy chain sequence of DH270min11, an antibody engineered from the DH270.6 bnAb to contain only those amino acid mutations determined to be functionally important for neutralization breadth [21]. Here **x** is the corresponding estimated UCA sequence for the DH270.6 clone [22] obtained by clonal lineage reconstruction using Clonalyst [10]. (Alternatively, a distribution over UCAs accounting for reconstruction uncertainty may be obtained by probabilistic methods such as Partis [23–25]; see Discussion.) In this case, **x** and **y** differ by seven nucleotides, and *a*(**x**) and *a*(**y**) differ by six amino acids. The DH270min11 mutations are given in Table 2. Notice that the largest separated set is of size *r*^*⋆*^ = 2 and *α* = 0.005 since *n* = 382. Of the observed amino acid mutations, only one requires multiple nucleotide substitutions in the corresponding codon.

**Table 2.**
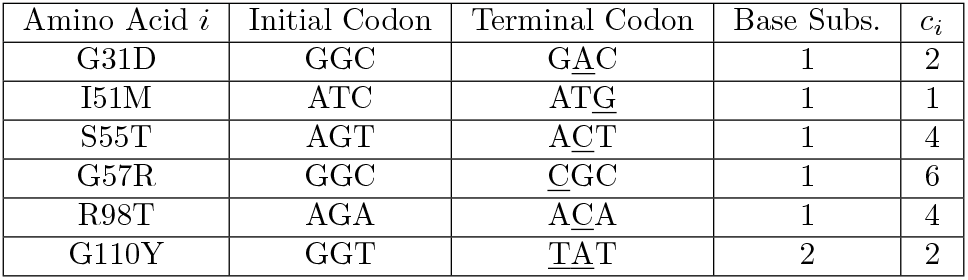
Description of the amino acid changes in the DH270min11 sequence. Columns show (in order): (1) amino acid change; (2) UCA sequence codon at change site; (3) corresponding codon observed in the DH270min11 sequence (mutations underlined); (4) minimal number of base substitutions required for amino acid change;(5) number of possible terminal codons that encode amino acid change.

We first consider the critical mutations individually in turn. So *q* = 1, and **m**_*i*_ consists of only a single amino acid location (i.e. we let **m**_*i*_ = {*m*_*i*_}, *i* = 1, …, *k*, and 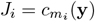, so **x**^*ij*^ is equal to **x** except at a single amino acid site. In total, there are 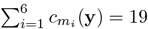 intermediate sequences **x**^*ij*^ corresponding to the total number of codons that encode the six amino acids where *a*(**x**) and *a*(**y**) differ. Table 3 lists the calculated path probabilities to DH270min11 conditional on each of the 6 individual amino acid mutations occurring first. To aid in the interpretability of results, we define an *inducibility index*

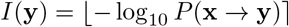

where ⌊⌉ denotes rounding to the nearest integer. This index equates sequences whose evolution probabilities are of the same order of magnitude, and facilitates direct comparison between potential targets with practically significant differences in inducibility. A lower inducibility index therefore indicates a sequence that has higher *a priori* probability of arising in the absence of selection.

**Table 3.** Transition probability results for *q* = 1. Initial amino acid mutations and corresponding codons are given in the first and second column. The starred codons correspond to the actual codons observed in the DH270min11 sequence. In the third column, we give the probability of transitioning to any nucleotide sequence **z** ∈ *A*(**y**) (i.e. any nucleotide sequence that gives rise to the same amino acid sequence as **y**) *conditional* on first obtaining the amino acid mutation **m**_*i*_ via codon *j*. In the fourth column, we give the estimate of the probability of obtaining the initial amino acid mutation in **m**_*i*_ by codon *j*. The weighted estimate in the fifth column is the product of the third and fourth columns. The sixth column is the average effective sample size of the transition probability estimates.

| $\mathbf{m}_i$ | Codon $j$ | $\hat{p}(A(\mathbf{y}) \mid \mathbf{x}^{ij})$ | $p(\mathbf{x}^{ij} \mid \mathbf{x}, r(\mathbf{x}^{ij}))$ | $\hat{p}(A(\mathbf{y}), \mathbf{x}^{ij} \mid \mathbf{x})$ | Avg. ESS | $I(\mathbf{y})$ |
| --- | --- | --- | --- | --- | --- | --- |
| **110Y** | **All** | **$3.29 \times 10^{-14}$** | $2.72 \times 10^{-6}$ | **$4.49 \times 10^{-20}$** | – | 13 |
| 110Y | TAC | $1.64 \times 10^{-14}$ | $1.30 \times 10^{-8}$ | $2.25 \times 10^{-22}$ | 8034 | 13 |
| 110Y | TAT* | $1.65 \times 10^{-14}$ | $2.71 \times 10^{-6}$ | $4.47 \times 10^{-20}$ | 8032 | 13 |
| **57R** | **All** | **$5.42 \times 10^{-15}$** | $1.30 \times 10^{-4}$ | **$1.24 \times 10^{-19}$** | – | 14 |
| 57R | AGA | $8.23 \times 10^{-16}$ | $8.19 \times 10^{-6}$ | $6.74 \times 10^{-21}$ | 7745 | 15 |
| 57R | AGG | $8.74 \times 10^{-16}$ | $4.37 \times 10^{-7}$ | $3.82 \times 10^{-22}$ | 7738 | 15 |
| 57R | CGA | $9.20 \times 10^{-16}$ | $2.89 \times 10^{-6}$ | $2.66 \times 10^{-21}$ | 7743 | 15 |
| 57R | CGC* | $9.63 \times 10^{-16}$ | $1.18 \times 10^{-4}$ | $1.14 \times 10^{-19}$ | 7746 | 15 |
| 57R | CGG | $9.82 \times 10^{-16}$ | $9.76 \times 10^{-8}$ | $9.59 \times 10^{-23}$ | 7746 | 15 |
| 57R | CGT | $8.58 \times 10^{-16}$ | $1.47 \times 10^{-7}$ | $1.26 \times 10^{-22}$ | 7743 | 15 |
| **98T** | **All** | **$1.27 \times 10^{-15}$** | $4.64 \times 10^{-4}$ | **$1.47 \times 10^{-19}$** | – | 14 |
| 98T | ACA* | $3.16 \times 10^{-16}$ | $4.62 \times 10^{-4}$ | $1.46 \times 10^{-19}$ | 7873 | 15 |
| 98T | ACC | $3.17 \times 10^{-16}$ | $9.60 \times 10^{-7}$ | $3.05 \times 10^{-22}$ | 7872 | 15 |
| 98T | ACG | $3.19 \times 10^{-16}$ | $3.73 \times 10^{-7}$ | $1.19 \times 10^{-22}$ | 7876 | 15 |
| 98T | ACT | $3.16 \times 10^{-16}$ | $5.30 \times 10^{-7}$ | $1.68 \times 10^{-22}$ | 7875 | 15 |
| **55T** | **All** | **$5.72 \times 10^{-16}$** | $1.02 \times 10^{-3}$ | **$1.47 \times 10^{-19}$** | – | 15 |
| 55T | ACA | $1.43 \times 10^{-16}$ | $5.56 \times 10^{-7}$ | $7.98 \times 10^{-23}$ | 7520 | 15 |
| 55T | ACC | $1.41 \times 10^{-16}$ | $1.10 \times 10^{-6}$ | $1.56 \times 10^{-22}$ | 7523 | 15 |
| 55T | ACG | $1.44 \times 10^{-16}$ | $4.49 \times 10^{-7}$ | $6.48 \times 10^{-23}$ | 7520 | 15 |
| 55T | ACT* | $1.44 \times 10^{-16}$ | $1.02 \times 10^{-3}$ | $1.47 \times 10^{-19}$ | 7527 | 15 |
| **51M** | **ATG\*** | **$2.94 \times 10^{-16}$** | $4.86 \times 10^{-4}$ | **$1.43 \times 10^{-19}$** | 7270 | 15 |
| **31D** | **All** | **$7.35 \times 10^{-17}$** | $3.94 \times 10^{-3}$ | **$1.47 \times 10^{-19}$** | – | 16 |
| 31D | GAC* | $3.74 \times 10^{-17}$ | $3.94 \times 10^{-3}$ | $1.47 \times 10^{-19}$ | 7566 | 16 |
| 31D | GAT | $3.61 \times 10^{-17}$ | $5.83 \times 10^{-6}$ | $2.11 \times 10^{-22}$ | 7573 | 16 |

We see that the transition probability 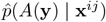 to the terminal DH270min11 sequence can be maximized by acquiring the G110Y mutation first. G110Y requires two base substitutions to occur within its codon for the amino acid transition from glycine (G) to tyrosine (Y). Comparing the calculated probabilities, we conclude that a vaccine regimen that successfully selects for the G110Y mutation first would increase the calculated probability of induction by 5 orders of magnitude, an order of magnitude (or more) improvement over selecting any of the other mutations first.

The vaccine designer can then use this information *(Design scenario I)* to aim for a priming immunogen that both binds with high affinity to the DH270 UCA (to initiate the clonal lineage), but also with even higher affinity to the UCA+G110Y mutation, in order to select for G110Y and guide the B cell response along the most probable bnAb maturation pathway.

As noted, the G110Y mutation requires two base changes. Multiple required base changes within a codon will typically result in a low transition probability of a targeted amino acid. However, it also provides an opportunity for the vaccine designer to accelerate its acquisition. For example, for the G110Y mutation, a single base change in codon 110 (GGT, glycine) of the DH270.6 UCA can transition through either GAT (aspartic acid) or TGT (cysteine). Our calculations indicate that the transition through aspartic acid is ≈ 1.5× more probable than the alternative path through cysteine.

Therefore, adding an immunogen to the vaccine regimen that selects for the intermediate amino acid state of aspartic acid at position 110 could accelerate induction of the critical and highly improbable G110Y mutation.

##### 3.2.3 Multiple simultaneous mutations in immunogen design calculations

In design scenarios 1 and 2, we may often wish to consider the induction of multiple simultaneous mutations. Here we consider initial target mutation sets of size *q* = 2 or *q* = 3. Tables 4 and 5 list the path probabilities conditional on all initial pairs and triplets of DH270min11 mutations. We see that different initial subsets differ by orders of magnitude in their probabilities, while the overall joint probabilities obtained as the products differ only by small constants (this also reassures that the approximation error is small in each case). We observe that (31D, 55T), (31D, 51M), and (31D, 98T) are the most probable pairs of mutations to arise in the absence of a selecting immunogen (have the highest “inducibility”), and (31D,51M,55T) and (31D,55T,98T) the most probable triples. Whereas (57R,110Y), (98T,110Y) and (51M,110Y) are the least probable pairs to arise by chance, and we therefore might expect them to be more difficult to elicit via immunogen selection. Similarly for the triples (57R,98T,110Y) and (51M,57R,110Y). Conversely, if we were able to design a priming immunogen to elicit the mutation pair (57R,110Y) or triplet (57R,98T,110Y), it would be of high impact as we would expect this to maximize the probability of obtaining the full mature bnAb using a boosting immunogen.

**Table 4.** Transition probability results for *q* = 2. The first columns indicates which amino acids are targeted first. Other columns are as described in Table 4.

| $\mathbf{m}_i$ | $\hat{p}(A(\mathbf{y}) \mid \mathbf{x}^{ij})$ | $\hat{p}(\mathbf{m}_i \mid \mathbf{x})$ | $\hat{p}(A(\mathbf{y}), \mathbf{m}_i \mid \mathbf{x})$ | Avg. Min. ESS | $I(\mathbf{y})$ |
| --- | --- | --- | --- | --- | --- |
| 57R, 110Y | $2.41 \times 10^{-10}$ | $1.11 \times 10^{-9}$ | $2.31 \times 10^{-20}$ | 7651 | 9 |
| 98T, 110Y | $5.18 \times 10^{-11}$ | $3.65 \times 10^{-9}$ | $2.37 \times 10^{-20}$ | 7641 | 10 |
| 55T, 110Y | $2.34 \times 10^{-11}$ | $8.05 \times 10^{-9}$ | $2.37 \times 10^{-20}$ | 7280 | 10 |
| 51M, 110Y | $1.21 \times 10^{-11}$ | $3.77 \times 10^{-9}$ | $2.29 \times 10^{-20}$ | 6794 | 10 |
| 57R, 98T | $7.28 \times 10^{-12}$ | $1.23 \times 10^{-7}$ | $3.87 \times 10^{-20}$ | 7457 | 11 |
| 55T, 57R | $3.28 \times 10^{-12}$ | $2.71 \times 10^{-7}$ | $3.88 \times 10^{-20}$ | 7121 | 11 |
| 31D, 110Y | $3.02 \times 10^{-12}$ | $3.10 \times 10^{-8}$ | $2.37 \times 10^{-20}$ | 7404 | 11 |
| 51M, 57R | $1.69 \times 10^{-12}$ | $1.28 \times 10^{-7}$ | $3.75 \times 10^{-20}$ | 6647 | 11 |
| 55T, 98T | $7.36 \times 10^{-13}$ | $9.30 \times 10^{-7}$ | $4.28 \times 10^{-20}$ | 7092 | 12 |
| 31D, 57R | $4.23 \times 10^{-13}$ | $1.04 \times 10^{-6}$ | $3.88 \times 10^{-20}$ | 7227 | 12 |
| 51M, 98T | $3.79 \times 10^{-13}$ | $4.38 \times 10^{-7}$ | $4.14 \times 10^{-20}$ | 6618 | 12 |
| 51M, 55T | $1.71 \times 10^{-13}$ | $9.66 \times 10^{-7}$ | $4.14 \times 10^{-20}$ | 6301 | 12 |
| 31D, 98T | $9.47 \times 10^{-14}$ | $3.58 \times 10^{-6}$ | $4.28 \times 10^{-20}$ | 7189 | 13 |
| 31D, 55T | $4.28 \times 10^{-14}$ | $7.90 \times 10^{-6}$ | $4.29 \times 10^{-20}$ | 6863 | 13 |
| 31D, 51M | $2.20 \times 10^{-14}$ | $3.72 \times 10^{-6}$ | $4.15 \times 10^{-20}$ | 6399 | 13 |

**Table 5.** Transition probability results for *q* = 3. The first column indicates which amino acids are targeted first. The second column corresponds to the probability of transitioning to **z** ∈ *A*(**y**) *conditional* on first obtaining the amino acid mutations in **m**_*i*_. The third column corresponds to the probability of *first* obtaining the amino acid mutations in **m**_*i*_. The fourth column corresponds to the probability of transitioning from **x** to any **z** ∈ *A*(**y**) (*not* the product of the second and third columns). The final column gives the average minimum effective sample size for the estimates of *p*(**z** | **x**^*ij*^, *r* = *d*_*H*_ (**x**^*ij*^, **z**)), where the minimum is across all **z** ∈ *Z*(**x, y**, *i, j*) and the average is across *j* = 1, …, *J*_*i*_.

| $\mathbf{m}_i$ | $\hat{p}(A(\mathbf{y}) \mid \mathbf{x}^{ij})$ | $\hat{p}(\mathbf{m}_i \mid \mathbf{x})$ | $\hat{p}(A(\mathbf{y}), \mathbf{m}_i \mid \mathbf{x})$ | Avg. Min. ESS | $I(\mathbf{y})$ |
| --- | --- | --- | --- | --- | --- |
| 57R, 98T, 110Y | $5.33 \times 10^{-7}$ | $2.14 \times 10^{-12}$ | $2.42 \times 10^{-20}$ | 8601 | 6 |
| 55T, 57R, 110Y | $2.41 \times 10^{-7}$ | $4.71 \times 10^{-12}$ | $2.42 \times 10^{-20}$ | 8210 | 6 |
| 51M, 57R, 110Y | $1.25 \times 10^{-7}$ | $2.19 \times 10^{-12}$ | $2.33 \times 10^{-20}$ | 7654 | 6 |
| 55T, 98T, 110Y | $4.95 \times 10^{-8}$ | $1.49 \times 10^{-11}$ | $2.31 \times 10^{-20}$ | 8205 | 7 |
| 31D, 57R, 110Y | $3.11 \times 10^{-8}$ | $1.81 \times 10^{-11}$ | $2.42 \times 10^{-20}$ | 8317 | 7 |
| 51M, 98T, 110Y | $2.57 \times 10^{-8}$ | $6.94 \times 10^{-12}$ | $2.23 \times 10^{-20}$ | 7653 | 7 |
| 51M, 55T, 110Y | $1.16 \times 10^{-8}$ | $1.53 \times 10^{-11}$ | $2.22 \times 10^{-20}$ | 7296 | 7 |
| 31D, 98T, 110Y | $6.39 \times 10^{-9}$ | $5.74 \times 10^{-11}$ | $2.31 \times 10^{-20}$ | 8312 | 8 |
| 55T, 57R, 98T | $5.76 \times 10^{-9}$ | $3.94 \times 10^{-10}$ | $2.43 \times 10^{-20}$ | 7992 | 8 |
| 51M, 57R, 98T | $2.98 \times 10^{-9}$ | $1.84 \times 10^{-10}$ | $2.35 \times 10^{-20}$ | 7455 | 8 |
| 31D, 55T, 110Y | $2.89 \times 10^{-9}$ | $1.26 \times 10^{-10}$ | $2.30 \times 10^{-20}$ | 7922 | 8 |
| 31D, 51M, 110Y | $1.50 \times 10^{-9}$ | $5.87 \times 10^{-11}$ | $2.22 \times 10^{-20}$ | 7405 | 8 |
| 51M, 55T, 57R | $1.34 \times 10^{-9}$ | $4.05 \times 10^{-10}$ | $2.35 \times 10^{-20}$ | 7118 | 8 |
| 31D, 57R, 98T | $7.42 \times 10^{-10}$ | $1.52 \times 10^{-9}$ | $2.43 \times 10^{-20}$ | 8097 | 9 |
| 31D, 55T, 57R | $3.35 \times 10^{-10}$ | $3.34 \times 10^{-9}$ | $2.43 \times 10^{-20}$ | 7723 | 9 |
| 51M, 55T, 98T | $2.89 \times 10^{-10}$ | $1.33 \times 10^{-9}$ | $2.41 \times 10^{-20}$ | 7088 | 9 |
| 31D, 51M, 57R | $1.73 \times 10^{-10}$ | $1.56 \times 10^{-9}$ | $2.35 \times 10^{-20}$ | 7224 | 9 |
| 31D, 55T, 98T | $7.21 \times 10^{-11}$ | $1.10 \times 10^{-8}$ | $2.50 \times 10^{-20}$ | 7713 | 10 |
| 31D, 51M, 98T | $3.72 \times 10^{-11}$ | $5.14 \times 10^{-9}$ | $2.41 \times 10^{-20}$ | 7195 | 10 |
| 31D, 51M, 55T | $1.68 \times 10^{-11}$ | $1.13 \times 10^{-8}$ | $2.41 \times 10^{-20}$ | 6858 | 10 |

##### 3.2.4 Comparison with experimental results

It is interesting to compare these results with recently obtained experimental data [20], where we have immunized DH270 UCA knock-in mice with a priming immunogen, sequenced their heavy chain BCR repertoires, and measured the frequency of the 6 DH270min11 mutations both individually and in combination. The G110Y mutation is observed to be the second least frequent mutation selected by our priming regimen. Our probability calculations (Table 3) indicate that of all individual mutations, G110Y selection maximizes bnAb maturation probability. Thus, adding a boosting immunogen to this vaccine regimen that can select for G110Y would be an optimal strategy for maximizing the probability of bnAb elicitation.

*(Design scenario III)* In our repertoire sequencing data, the I51M mutation is the lowest frequency of the six DH270min11 mutations. Contrasting this with our probability calculations (Table 3),which estimate that I51M is the third most probable mutation in the absence of selection. Such differences between model calculations and observed frequencies may be indicative of selection effects; thus one explanation for the low I51M frequency observed in immunized mice is that our priming immunogen lacks sufficient selection strength for this mutation and may even be selecting against it.

##### 3.2.5 Design scenario II

When information is available about the performance of the first immunogen(s) in a sequential boost vaccine regimen, the vaccine designer can use the estimated transition probabilities to make decisions about which boosting immunogen(s) to administer next in the series. From our experimental data, the highest frequency mutation pair induced by our priming immunogen was (G31D, R98T).

For current purposes, we assume a single immunogen can select for only one pair of 313 mutations, and use the model calculations to choose which pair of mutations should be 314 selected for with the next boosting immunogen. Table 6 shows the estimated transition 315 probabilities starting from the UCA + (G31D,R98T), for all remaining 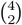 pairs of 316 mutations. We observe that the transition probability to the DH270min11 sequence is 317 maximized upon acquiring (57R,110Y). Thus, the optimal sequential boosting strategy 318 is to use a first boosting immunogen to select for G57R and G110Y followed by a 319 second boosting immunogen to select for S55T and I51M.

**Table 6.** Starting from (G31D, R98T), the most frequent pair observed in immunized mice, and transitioning to the next pair.

| $\mathbf{m}_i$ | $\hat{p}(A(\mathbf{y}) \mid \mathbf{x}_{i,j})$ | $\hat{p}(\mathbf{m}_i \mid \mathbf{x})$ | $\hat{p}(A(\mathbf{y}), \mathbf{m}_i \mid \mathbf{x})$ | Avg. Min.<br>ESS | $I(\mathbf{y})$ |
| --- | --- | --- | --- | --- | --- |
| 57R,110Y | $1.15 \times 10^{-5}$ | $1.13 \times 10^{-9}$ | $1.10 \times 10^{-15}$ | 9334 | 4 |
| 55T,110Y | $1.02 \times 10^{-6}$ | $8.20 \times 10^{-9}$ | $1.05 \times 10^{-15}$ | 8903 | 5 |
| 51M,110Y | $5.31 \times 10^{-7}$ | $3.84 \times 10^{-9}$ | $1.02 \times 10^{-15}$ | 8319 | 6 |
| 55T,57R | $9.25 \times 10^{-8}$ | $2.75 \times 10^{-7}$ | $1.09 \times 10^{-15}$ | 8670 | 7 |
| 51M,57R | $4.80 \times 10^{-8}$ | $1.29 \times 10^{-7}$ | $1.06 \times 10^{-15}$ | 8085 | 7 |
| 51M,55T | $4.46 \times 10^{-9}$ | $9.78 \times 10^{-7}$ | $1.09 \times 10^{-15}$ | 7735 | 8 |

### 4 Conclusion

We have introduced a model-based approach to sequential immunogen design using the ARMADiLLO model of somatic hypermutation. To calculate design-relevant marginal sequence transition probabilities in the face of context-dependent mutation, we have developed a fast and accurate Monte Carlo approximation scheme. We have demonstrated that this model performs well on test sequences of varying complexity. Finally, we have applied this approach to answer questions of great current significance regarding mutation targeting for boosting immunogen design for ongoing efforts to elicit the HIV bnAb DH270min11. These results are now being used to guide efforts for immunogen design in the Duke Human Vaccine Institute.

Our approach relies on knowledge of a precursor sequence. Typically this is obtained as an estimated UCA obtained from a set of clonally related sequences. However, the process of inferring the UCA retains residual uncertainty both due to choices regarding which sequences to include, and the probabilistic information content of the sequences themselves. Although we use the Clonalyst procedure here to produce a single maximum likelihood UCA, other methods [23] are available which account for (some of) the reconstruction uncertainty by providing a *distribution* over UCA sequences, and our approach easily extends to use such information. However those methods rely on phylogenetic computations which become intractable in the face of context-dependent mutation, and so do not currently account for this aspect of the somatic hypermutation process, which has been our focus here. Applying the accurate, rapid approximation of marginal sequence likelihoods developed here to the problem of lineage reconstruction in the face of context-dependent mutation is a promising area for future work. We expect that this may only impact results in the CDRH3 region, as the posterior probabilities on V genes and even alleles are expected to be close to one, but it may indeed be important for the CDRH3 (R98T and G110Y are in the CDRH3, for example). Another source of uncertainty in the inferred UCA arises from the selection of sequences to define the clone itself; we are exploring approaches to account for context-dependence in this step as well. Finally, given the speed of computation, our approach here could be extended to sets of bnAb UCAs that define an entire precursor class, i.e. a common set of precursor sequences evolving to a common set of paratopic features that define that bnAb class’s ability to recognize a conserved site of vulnerability on a pathogen.

## Acknowledgements

This work was supported by NIH Division of AIDS grant UM1AI144371 for the Duke Consortia for HIV/AIDS Vaccine Development (KW and JM). The content is solely the responsibility of the authors and does not necessarily represent the official views of the National Institutes of Health.

## References

1. Walker LM, Burton DR. Passive Immunotherapy of Viral Infections: ‘Super-Antibodies’ Enter the Fray. Nat Rev Immunol. 2018;18(5):297–308. doi:10.1038/nri.2017.148.

2. Haynes BF, Wiehe K, Borrrow P, Saunders KO, Korber B, Wagh K, et al. Strategies for HIV-1 Vaccines that Induce Broadly Neutralizing Antibodies. Nat Rev Immunol. 2022;doi:10.1038/s41577-022-00753-w.

3. Jardine JG, Kulp DW, Havenar-Daughton C, Sarkar A, Briney B, Sok D, et al. HIV-1 Broadly Neutralizing Antibody Precursor B Cells Revealed by Germline-Targeting Immunogen. Science. 2016;351(6280):1458–63. doi:10.1126/science.aad9195.

4. Wiehe K, Bradley T, Meyerhoff RR, Hart C, Williams WB, Easterhoff D, et al. Functional Relevance of Improbable Antibody Mutations for HIV Broadly Neutralizing Antibody Development. Cell Host Microbe. 2018;23(6):759–765 e6. doi:10.1016/j.chom.2018.04.018.

5. Saunders KO, Wiehe K, Tian M, Acharya P, Bradley T, Alam SM, et al. Targeted Selection of HIV-Specific Antibody Mutations by Engineering B Cell Maturation. Science. 2019;366(6470). doi:10.1126/science.aay7199.

6. Haynes BF, Kelsoe G, Harrison SC, Kepler TB. B-Cell-Lineage Immunogen Design in Vaccine Development with HIV-1 as a Case Study. Nat Biotechnol. 2012;30(5):423–33. doi:10.1038/nbt.2197.

7. Jardine JG, Ota T, Sok D, Pauthner M, Kulp DW, Kalyuzhniy O, et al. HIV-1 VACCINES. Priming a Broadly Neutralizing Antibody Response to HIV-1 Using a Germline-Targeting Immunogen. Science. 2015;349(6244):156–61. doi:10.1126/science.aac5894.

8. Steichen JM, Lin YC, Havenar-Daughton C, Pecetta S, Ozorowski G, Willis JR, et al. A Generalized HIV Vaccine Design Strategy for Priming of Broadly Neutralizing Antibody Responses. Science. 2019;366(6470). doi:10.1126/science.aax4380.

9. Kwong PD, Mascola JR. HIV-1 Vaccines Based on Antibody Identification, B Cell Ontogeny, and Epitope Structure. Immunity. 2018;48(5):855–871. doi:10.1016/j.immuni.2018.04.029.

10. Kepler TB. Reconstructing a B-Cell Clonal Lineage. I. Statistical Inference of Unobserved Ancestors. F1000Res. 2013;2:103. doi:10.12688/f1000research.2-103.v1.

11. Leggat DJ, Cohen KW, Willis JR, Fulp WJ, deCamp AC, Kalyuzhniy O, et al. Vaccination Induces HIV Broadly Neutralizing Antibody Precursors in Humans. Science. 2022;378(6623):eadd6502. doi:10.1126/science.add6502.

12. Caniels TG, Medina-Ramirez M, Zhang J, Sarkar A, Kumar S, LaBranche A, et al. Germline-Targeting HIV-1 Env Vaccination Induces VRC01-Class Antibodies with Rare Insertions. Cell Rep Med. 2023;4(4):101003. doi:10.1016/j.xcrm.2023.101003.

13. Tian M, Cheng C, Chen X, Duan H, Cheng HL, Dao M, et al. Induction of HIV Neutralizing Antibody Lineages in Mice with Diverse Precursor Repertoires. Cell. 2016;166(6):1471–1484 e18. doi:10.1016/j.cell.2016.07.029.

14. Briney B, Sok D, Jardine JG, Kulp DW, Skog P, Menis S, et al. Tailored Immunogens Direct Affinity Maturation Toward HIV Neutralizing Antibodies. Cell. 2016;166(6):1459–1470 e11. doi:10.1016/j.cell.2016.08.005.

15. Yaari G, Vander Heiden JA, Uduman M, Gadala-Maria D, Gupta N, Stern JN, et al. Models of Somatic Hypermutation Targeting and Substitution Based on Synonymous Mutations from High-Throughput Immunoglobulin Sequencing Data. Front Immunol. 2013;4:358. doi:10.3389/fimmu.2013.00358.

16. Jensen J, Pederson AM. Probabilistic Models of DNA Sequence Evolution with Context Dependent Rates of Substitution. Advances in Applied Probability. 2000;32(2):499–517.

17. Hobolth A. A Markov Chain Monte Carlo Expectation Maximization Algorithm for Statistical Analysis of DNA Sequence Evolution with Neighbor-Dependent Substitution Rates. Journal of Computational and Graphical Statistics. 2008;17(1):138–162. doi:10.1198/106186008x289010.

18. Robinson DM, Jones DT, Kishino H, Goldman N, Thorne JL. Protein Evolution with Dependence Among Codons due to Tertiary Structure. Mol Biol Evol. 2003;20(10):1692–704. doi:10.1093/molbev/msg184.

19. Felenstein J. Evolutionary Trees from DNA Sequences: A Maximum Likelihood Approach. Journal of Molecular Evolution. 1981;17:368–376.

20. Wiehe K, Saunders KO, Stalls V, Cain DW, Venkatayogi S, Beem JSM, et al. Mutation-Guided Vaccine Design: A Strategy for Developing Boosting Immunogens for HIV Broadly Neutralizing Antibody Induction. bioRxiv. 2022;doi:10.1101/2022.11.11.516143.

21. Swanson O, Rhodes B, Wang A, Xia SM, Parks R, Chen H, et al. Rapid Selection of HIV Envelopes that Bind to Neutralizing Antibody B Cell Lineage Members with Functional Improbable Mutations. Cell Rep. 2021;36(7):109561. doi:10.1016/j.celrep.2021.109561.

22. Bonsignori M, Kreider EF, Fera D, Meyerhoff RR, Bradley T, Wiehe K, et al. Staged Induction of HIV-1 Glycan-Dependent Broadly Neutralizing Antibodies. Sci Transl Med. 2017;9(381). doi:10.1126/scitranslmed.aai7514.

23. Ralph DK, Matsen FA. Consistency of VDJ Rearrangement and Substitution Parameters Enables Accurate B Cell Receptor Sequence Annotation. PLoS Comput Biol. 2016;12(1):e1004409. doi:10.1371/journal.pcbi.1004409.

24. Ralph DK, Matsen FA. Likelihood-Based Inference of B Cell Clonal Families. PLoS Comput Biol. 2016;12(10):e1005086. doi:10.1371/journal.pcbi.1005086.

25. Dhar A, Ralph DK, Minin VN, Matsen FA. A Bayesian Phylogenetic Hidden Markov Model for B Cell Receptor Sequence Analysis. PLoS Comput Biol. 2020;16(8):e1008030. doi:10.1371/journal.pcbi.1008030.

